# The Multispecies Coalescent in Space and Time

**DOI:** 10.1101/2020.08.02.233395

**Authors:** Patrick F. McKenzie, Deren A. R. Eaton

## Abstract

A key distinction between species tree inference under the multi-species coalescent model (MSC), and the inference of gene trees in sliding windows along a genome, is in the effect of genetic linkage. Whereas the MSC explicitly assumes genealogies to be unlinked, i.e., statistically independent, genealogies located close together on genomes are spatially auto-correlated. Here we use tree sequence simulations with recombination to explore the effects of species tree parameters on spatial patterns of linkage among genealogies. We decompose coalescent time units to demonstrate differential effects of generation time and effective population size on spatial coalescent patterns, and we define a new metric, “phylogenetic linkage,” for measuring the rate of decay of phylogenetic similarity by comparison to distances among unlinked genealogies. Finally, we provide a simple example where accounting for phylogenetic linkage in sliding window analyses improves local gene tree inference.

## 0.1 Introduction

The multispecies coalescent (MSC) is a model for inferring a phylogenetic tree from a distribution of sampled genealogies – or in practice, a distribution of empirical gene trees inferred from multi-locus genetic data (Maddison, 1997; Maddison & Knowles, 2006; Degnan & Rosenberg, 2009). By integrating over genealogical variation the MSC improves estimation of both tree topologies and divergence times, in addition to providing estimates of other demographic parameters of interest, such as population sizes (Edwards & Beerli, 2000; Fang *et al.*, 2020). Its influence on phylogenetics has been broad and pervasive, as is evident in the many extensions that have been developed for incorporating the MSC into studies of gene duplication and loss (Rasmussen & Kellis, 2012), introgression (Yu *et al.*, 2011), and even character evolution (Guerrero & Hahn, 2018). As we approach one decade since the publication of the first volume of *Estimating Species Trees* (Knowles & Kubatko, 2011) it is valuable to re-examine the MSC, and its assumptions, to ask how we can best approach new challenges and opportunities in the coming era of ubiquitous whole genome data sets. One area where we believe the MSC has great potential is in improving the inference of local genealogical variation across whole genomes.

A key distinction between species tree inference under the MSC and the inference of genealogies sequentially distributed across genomes is the effect of genetic linkage. The MSC explicitly assumes that genealogies are unlinked, i.e., statistically independent, whereas genealogies distributed across a contiguous genomic region are not independent, and are expected to be spatially auto-correlated. This correlation (linkage disequilibrium) decays over time as recombination causes samples within different genomic regions to trace back to different sampled ancestors (Hudson & Kaplan, 1988). While this decay function has been well studied in the context of single populations (McVean & Cardin, 2005), its effect on the similarity of genealogies constrained by a species tree model is poorly understood, including the influence of species tree parameters. Recent algorithmic advances have now made it possible to efficiently simulate entire chromosomes with recombination to produce correlated tree sequences (Kelleher *et al.*, 2016), which presents a powerful new opportunity to investigate the relationship between species tree parameters and sequential genealogical patterns across genomes.

Genome-wide phylogenetic inference is currently approached from two methodological extremes: either (1) a single species tree is inferred as a hierarchical model to describe the expected distribution of unlinked genealogies across the genome; or (2) no hierarchical model is assumed, and gene trees are inferred independently in sliding windows of concatenated sequences along the genome (Martin & Belleghem, 2017). The latter approach is often applied to identify introgressed regions based on their deviation from a genome-wide average (Wang *et al.*, 2019). However, the dearth of information contained within small genomic windows can cause high gene tree estimation error in this approach, and similarly, increasing window size to be too large will cause errors from concatenation of multiple distinct histories. The MSC provides a potential path forward. A parameterized species tree inferred from unlinked locus data may be able to provide priors on the expected distribution of genealogies both globally across the genome, as well as spatially among linked trees.

In this chapter we explore this concept by using simulations to estimate the effect of species tree parameters on the rate of decay of phylogenetic similarity across the spatial extent of a chromosome with a uniform recombination rate. We show that a decay function can be estimated to describe the spatial auto-correlation of genealogies, and that by incorporating this function into gene tree inference, accuracy can be significantly improved compared to existing sliding window methods.

## 0.2 Coalescent simulations

To investigate genealogical variation along chromosomes we simulated genealogies under a range of species tree models in Python using the *ipcoal* package (McKenzie & Eaton, 2020). This takes as input a tree topology and demographic parameters (divergence times, effective population sizes, mutation rate, and recombination rate) to generate a parameterized simulator for the program *msprime* (Kelleher *et al.*, 2016). Using this model we then simulated coalescent genealogies constrained by a species tree topology. To generate linked trees we simulated a 1Mb chromosome and recorded the true genealogy spanning each position of its length, since different genealogies span different intervals along the chromosome between recombination crossover locations. To generate unlinked trees we simulated 1000 independent loci of length one and stored the single observed genealogy from each locus. Species trees and genealogies were plotted and manipulated using the Python package *toytree* (Eaton, 2020). Annotated code to reproduce all analyses in this chapter is organized into jupyter notebooks and available at https://github.com/eaton-lab/sptree-chapter.

The distributions of linked and unlinked genealogies simulated on the same species tree are easy to distinguish when visualized: linked genealogies exhibit significant auto-correlation whereas unlinked genealogies exhibit greater variation (Fig. 1c-d). We explored a range of parameters to realistically describe linked and unlinked genealogical variation in genome-wide phylogenetic data sets. To focus our analyses on fewer total parameters we performed all simulations on completely imbalanced tree shapes (but different tree sizes) in which internode lengths and effective population sizes of internal edges are all set to be equal. All simulations were performed using a per-site per-generation recombination rate of 1e-9, and in the case when sequence data was generated, a per-site per-generation mutation rate of 1e-8 applied under the JC69 substitution model. The parameters we investigated for their effect on the distribution of genealogies include tree size (number of tips), the probability of incongruence (internode edge lengths in coalescent units), and tree height (the number of generations between internodes).

**Figure 1.**
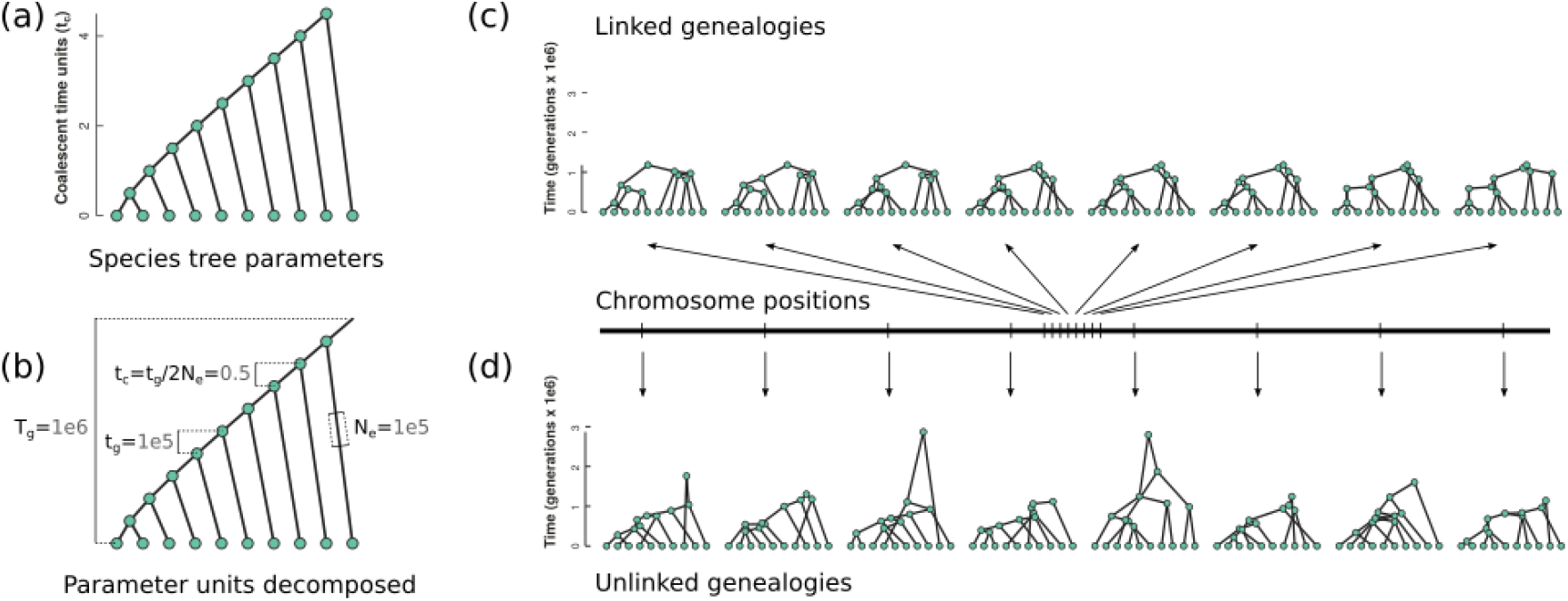
The effect of genetic linkage on the spatial distribution of genealogies across a chromosome. (a) A species tree topology with edge lengths in coalescent units can fully describe the probability of incongruence among unlinked genealogies. (b) Coalescent units (t_*c*_) are a composite of time in generations (t_*g*_) and the effective populations size (N_*e*_). The extent of linkage among genealogies is influenced by both t_*g*_ and N_*e*_, and thus not fully explained by t_*c*_ alone. (c-d) Genealogies are plotted with tips in the same order as the species tree topology to highlight incongruence. Arrows indicate the positions of genealogies on a chromosome; linked genealogies are close together and unlinked genealogies are far apart. (c) Linked genealogies are spatially correlated because many samples share the same ancestors until a recombination event occurs. (d) Unlinked genealogies are independent and exhibit greater variation among a sampled set than linked genealogies.

### 0.2.1 Units, space, and time

The effect of time, measured in units of generations, is not typically of interest for studies of the MSC, since the probability of incongruence (among unlinked genealogies) can be explained entirely by internode lengths measured in coalescent units (*t*_*c*_), which is calculated as *t*_*c*_ = *t*_*g*_/2*N*_*e*_, where *t*_*g*_ is time in generations and *N*_*e*_ is the effective population size. Because *t*_*c*_ is a ratio of time and population size, the absolute value of *t*_*g*_ has not been of interest, only its relation to *N*_*e*_ (Fig. 1a-b). However, in the context of a sequential coalescent process it turns out that *t*_*g*_ does matter, since recombination is modeled as a per-generation process, and so both *t*_*g*_ and *N*_*e*_ affect the number of recombination events, and thus the similarity of neighboring genealogies.

The effect of time in units of generations is demonstrated in Table 1. Here we simulated linked and unlinked genealogies on the same species trees and over a range of parameters. In each data set we measured the average Robinson-Foulds (RF) distance between all pairwise unlinked genealogies, and in the case of linked trees, between 1000 pairs of genealogies randomly sampled from positions that are spaced 5Kb apart. RF distances are reported here using normalized (scaled) values to account for differences in tree size, but non-normalized RF distances show the same qualitative results (not shown). In the unlinked data sets *t*_*g*_ has no effect on the similarity of genealogies – only *t*_*c*_ is relevant – as has been traditionally recognized in the MSC. However, for linked genealogies *t*_*g*_ has a large effect. When edge lengths are longer in units of generations, and N_*e*_ is similarly scaled to retain the same probability of incongruence (t_*c*_), the size of non-recombined blocks becomes smaller, and the average RF distance between neighboring genealogies is greater.

**Table 1.** Parameter settings used in simulations to examine the distribution of linked versus unlinked genealogies generated on the same species tree. All simulations were performed on an imbalanced species tree with uniform internode edge lengths. Three free parameters were explored: the number of tips (Ntips) on the tree, total tree height in generations (T_*g*_), and internode edge lengths in coalescent units (t_*c*_). Two additional parameters are shown for which values were determined entirely by values of the free parameters: the internode length in units of generations (t_*g*_) is determined by T_*g*_ and Ntips, and effective population size (N_*e*_) is determined by t_*c*_ and t_*g*_. Results are reported as the mean values calculated from 1000 simulated genealogies. The size of non-recombined genomic blocks (block-size) decreases with time in generations. This affects the RF distance between linked genealogies, but not unlinked genealogies. RF_*linked*−5*K*_ is the RF distances among linked trees separated by 5Kb on a chromosome. The phylogenetic half-life was calculated from fitting an exponential curve to the rate of decay of phylogenetic linkage.

| | Ntips | $T_g$ | $t_c$ | $t_g$ | $N_e$ | block-size | $RF_{unlinked}$ | $RF_{linked-5K}$ | half-life |
| --- | --- | --- | --- | --- | --- | --- | --- | --- | --- |
| 0 | 10 | 1e5 | 0.2 | 1e4 | 25000 | 1706 | 0.40 | 0.19 | 6576 |
| 1 | 10 | 1e6 | 0.2 | 1e5 | 250000 | 174 | 0.40 | 0.40 | 607 |
| 2 | 10 | 1e7 | 0.2 | 1e6 | 2500000 | 18 | 0.40 | 0.40 | 67 |
| 3 | 10 | 1e5 | 1.0 | 1e4 | 5000 | 4761 | 0.22 | 0.05 | 16715 |
| 4 | 10 | 1e6 | 1.0 | 1e5 | 50000 | 525 | 0.22 | 0.15 | 2359 |
| 5 | 10 | 1e7 | 1.0 | 1e6 | 500000 | 56 | 0.22 | 0.20 | 270 |
| 6 | 10 | 1e5 | 2.0 | 1e4 | 2500 | 8849 | 0.08 | 0.01 | 22627 |
| 7 | 10 | 1e6 | 2.0 | 1e5 | 25000 | 906 | 0.08 | 0.05 | 4659 |
| 8 | 10 | 1e7 | 2.0 | 1e6 | 250000 | 93 | 0.08 | 0.08 | 443 |
| 9 | 50 | 1e5 | 0.2 | 2e3 | 5000 | 1485 | 0.47 | 0.13 | 15890 |
| 10 | 50 | 1e6 | 0.2 | 2e4 | 50000 | 149 | 0.47 | 0.39 | 1838 |
| 11 | 50 | 1e7 | 0.2 | 2e5 | 500000 | 15 | 0.47 | 0.47 | 185 |
| 12 | 50 | 1e5 | 1.0 | 2e3 | 1000 | 4830 | 0.24 | 0.01 | 84731 |
| 13 | 50 | 1e6 | 1.0 | 2e4 | 10000 | 509 | 0.24 | 0.09 | 11884 |
| 14 | 50 | 1e7 | 1.0 | 2e5 | 100000 | 51 | 0.24 | 0.22 | 1233 |
| 15 | 50 | 1e5 | 2.0 | 2e3 | 500 | 9900 | 0.09 | 0.00 | 124415 |
| 16 | 50 | 1e6 | 2.0 | 2e4 | 5000 | 866 | 0.09 | 0.02 | 23019 |
| 17 | 50 | 1e7 | 2.0 | 2e5 | 50000 | 87 | 0.09 | 0.08 | 2048 |
| 18 | 100 | 1e5 | 0.2 | 1e3 | 2500 | 1345 | 0.47 | 0.08 | 26873 |
| 19 | 100 | 1e6 | 0.2 | 1e4 | 25000 | 138 | 0.47 | 0.34 | 3128 |
| 20 | 100 | 1e7 | 0.2 | 1e5 | 250000 | 14 | 0.47 | 0.47 | 345 |
| 21 | 100 | 1e5 | 1.0 | 1e3 | 500 | 5405 | 0.24 | 0.01 | 178444 |
| 22 | 100 | 1e6 | 1.0 | 1e4 | 5000 | 474 | 0.24 | 0.05 | 22688 |
| 23 | 100 | 1e7 | 1.0 | 1e5 | 50000 | 51 | 0.24 | 0.19 | 2499 |
| 24 | 100 | 1e5 | 2.0 | 1e3 | 250 | 7299 | 0.09 | 0.00 | 218558 |
| 25 | 100 | 1e6 | 2.0 | 1e4 | 2500 | 877 | 0.09 | 0.01 | 39616 |
| 26 | 100 | 1e7 | 2.0 | 1e5 | 25000 | 85 | 0.09 | 0.06 | 4169 |

A notable result of these simulations is the observation that the size of non-recombined genomic blocks becomes very small in certain regions of parameter space, particularly when the internode lengths in units of generations are very long. This is troubling for the MSC which requires that loci represent a single genealogical history as opposed to multiple concatenated genealogies. The impact of recombination within loci has been investigated previously, both in the first edition of this book (Castillo-Ramirez *et al.*, 2010), as well as in a series of critical examinations of the impact of “concatalescence” (Springer & Gatesy, 2016). At issue is whether gene tree estimation error is elevated when concatenating data from multiple genealogies into a single locus. The results from our simulations suggest there are some regions of parameter space where the size of non-recombined blocks becomes quite small, and so this issue may warrant further examination. In this paper, however, rather than examine recombination and its effects as a critique on existing MSC approaches, we aim instead to explore how linkage among recombined genealogical blocks of the genome can possibly be a useful source of information when analyzing whole genomes.

### 0.2.2 Tree size, tree space, and phylogenetic decay

The enormous size of phylogenetic tree space is a constant source of computational burden in phylogenetics, but intriguingly, it may actually provide a source of information in the context of the sequential coalescent process. This is because as the size of tree space grows in larger data sets so too does the expected RF distance between any two random unlinked genealogies. This is particularly true when *t*_*c*_ is very small, such that all coalescent events occur deeper than the root of the species tree. In this case the topology of unlinked genealogies is hardly constrained by the species tree at all, and almost any genealogy can be observed. However, adjacent genealogies on the same chromosome are still expected to share significant similarity, since few recombination events are likely to have occurred between them. Consequently, the degree to which linked genealogies are more similar to each other, *relative* to the similarity among unlinked genealogies, is a function of parameters of the species tree, including the tree size.

This type of relative measurement provides a means to develop a statistic to describe the rate of decay of spatial auto-correlation in genealogies across a genome. We propose the term “phylogenetic linkage” (*PL*) to describe the ratio of RF distances among linked genealogies separated by some genetic distance in the genome relative to the average RF distance among unlinked genealogies.

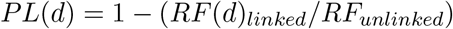

In other words, if two genealogies spaced *d* distance apart on a chromosome are as different from each other as two randomly sampled unlinked genealogies are on average, then they are effectively unlinked. By measuring phylogenetic linkage at increasing genetic distances between genealogies we can infer a rate of decay of phylogenetic linkage across the genome. For each simulated data set we then fit an exponential decay function using the *scipy* package in Python. From the estimated decay rate parameter (*λ*_*d*_) we estimated a phylogenetic linkage half-life, representing the distance in bp at which two genealogies are expected to lose half of their phylogenetic linkage (Table 1; Fig. 2).

**Figure 2.**
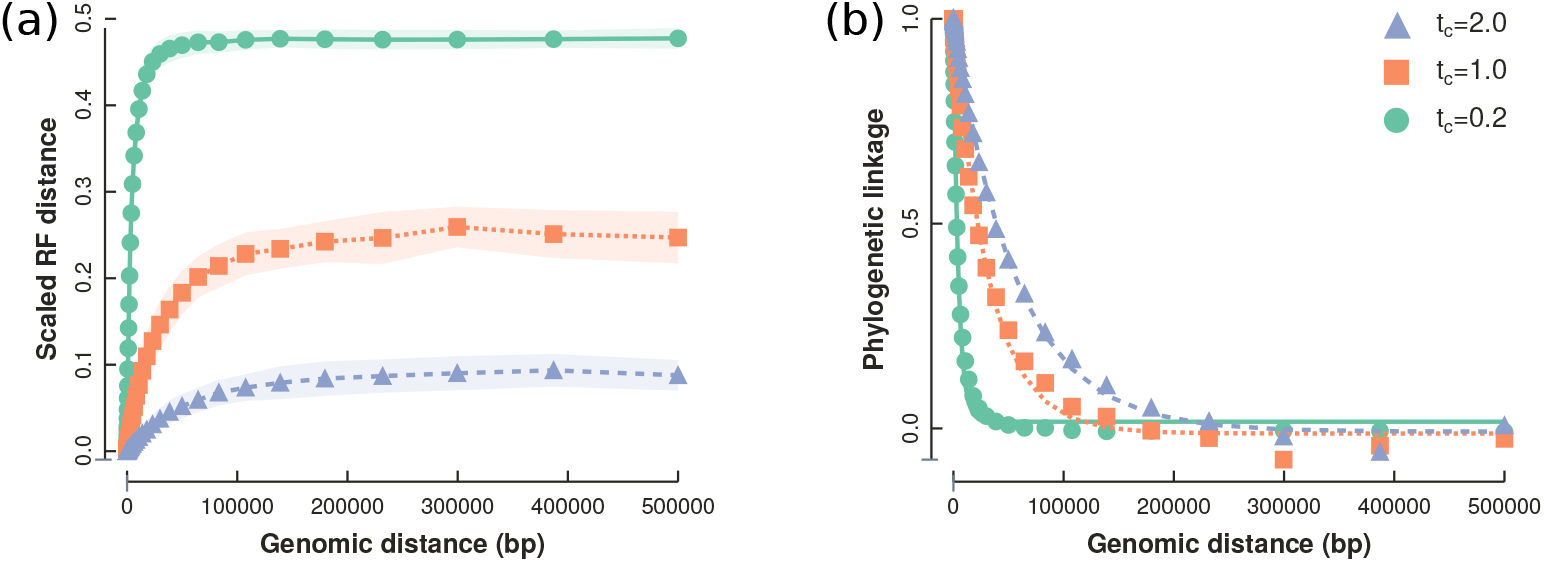
The RF distance between genealogies separated spatially on a chromosome plateaus as linkage decays by recombination, and approaches the average RF distance between unlinked genealogies (a). The ratio of RF distances between linked and unlinked genealogies (phylogenetic linkage) measured at different genetic distances approximates an exponential decay function (b). Results are shown for data simulated on a 100 tip species tree with total tree height of *T*_*g*_=1e6 generations, and *N*_*e*_ of 2.5e4, 5e4, or 2.5e3, corresponding to edge lengths in coalescent units (*t*_*c*_) of 0.2, 1.0, and 2.0, respectively.

When t_*g*_ is larger, phylogenetic decay occurs faster, since more recombination events are possible over each internal edge of the tree (e.g., compare rows 0 and 7 in Table 1). Similarly, when *N*_*e*_ is greater, recombination events are more likely to cause a change in the topology, and thus phylogenetic linkage decays faster (e.g., compare rows 0, 3, and 6 in Table 1). Finally, when the total tree size (Ntips) is greater, the decay of linkage occurs more slowly, since the average difference between unlinked genealogies is greater, and thus it takes longer for sufficient spatial information to decay to approach the unlinked mean RF distance (e.g., compare rows 0, 9, and 18 in Table 1).

## 0.3 Linked genealogies and gene tree inference

Unlike in simulations, the true genealogical history for any region of the genome is an unknown and unobservable variable. It is something we must infer based on the signal left by the mutational process. So how can our understanding of the decay of phylogenetic linkage be useful in the context of gene tree inference? One way to approach this problem is to ask what is the expected length over which a site supporting a bipartition in one position of the genome continues to be true in neighboring regions of the genome?

The standard sliding window approach gives equal weight to all sites within an alignment window. An alternative approach could be to extend the size of the window to ensure that there is sufficient information to infer a resolved gene tree, but to apply variable weights to sites in the alignment such that those near the center of the window have greatest weight, and can override alternative signals (Fig. 3a). This has the effect that if no local information exists to support the true local genealogy then data from more distant regions can inform that part of the tree, but with decreasing weight as their probability of representing the same genealogy decays with distance.

**Figure 3.**
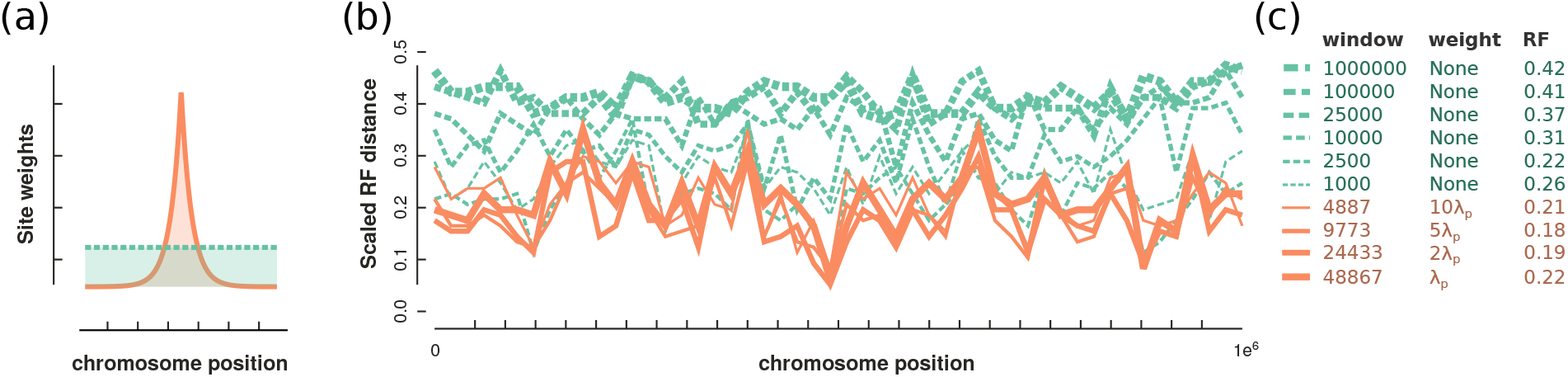
The accuracy of gene tree inference in sliding windows along a chromosome. (a) We compared windows that were of a fixed length and with uniform site weights to windows with lengths and weights determined by a function of exponentially decaying phylogenetic linkage inferred from simulations under the species tree parameters. (b) The scaled RF distance between inferred gene trees and the true simulated genealogy at 50 positions across a genome measured with different window sizes and weightings. (c) Windows with uniform (no) weights inferred less accurate gene trees than those with weights distributed by a phylogenetic decay-function, as measured by the mean scaled RF distance to the true simulated genealogies.

We implemented a weighted approach to local gene tree inference by using the “-a” weights file argument during maximum likelihood tree estimation in RAxML v.8.2.12 (Stamatakis, 2014). A distribution of site weights was generated to give exponentially decreasing weight to sites on either side of a central position. For computational efficiency we cut off the window size to the left and right of the center at sites where the weight reached 1/10000 of the center given the exponential decay rate parameter estimated for that data set.

Decay-weighted gene tree inference was compared to traditional windows with uniform (no) weights. Gene trees were inferred at 50 positions spaced evenly across a 1Mb simulated chromosome. At each position the RF distance between the true genealogy and the inferred gene tree was recorded to measure gene tree estimation accuracy. We tested uniform windows of lengths 1Kb, 2.5Kb, 10Kb, 25Kb, 100Kb, and 1Mb, the last of which represents the total concatenated chromosome gene tree. Decay-function weighted gene trees were estimated in windows with a size determined by the decay rate, and we additionally tested decay rates with 2X, 5X and 10X faster rates to examine sensitivity to rate estimation. We show results for simulations performed on data set 10 from Table 1, which was selected for its fast rate of decay and high incongruence so that many distinct genealogies would be observed across the chromosome.

The decay-function weighted windows inferred more accurate gene trees on average than uniform windows (Fig. 3b-c). Of the uniform windows, the largest size (representing concatenation of the entire chromosome) performed the worst, while the best window size appears to be near 2.5Kb. All four decay-function weighted window sizes tested had lower mean RF scores (greater accuracy) than the best scoring uniform window. The best estimate was observed for the 5X decay rate window, which had a mean scaled RF distance of only 0.18, making it more than twice as accurate as the genome-wide concatenation gene tree. In non-scaled RF scores this represents an average of 35 differences from the true genealogy compared to 43 differences in the 2.5Kb uniform windows, 51 in 10Kb windows, and 80 in the concatenation gene tree. The reason a 5X decay rate performed better than the estimated decay rate may be caused by the cut off to weighted window sizes that we implemented to improve run times. Additional parameters that we did not explore here, such as the mutation rate and recombination rate, are likely to be important factors as well, since they affect the information content within each non-recombined genomic block.

## Conclusions

For over a decade the goal of phylogenomic analyses has primarily focused on inferring a single species tree to represent the distribution of genealogical variation across the entire genome. However, as whole genome data becomes available there is increasing interest in the spatial distribution of genealogies at specific locations across the genome. This type of local ancestry information can be useful for testing evolutionary questions about patterns of hemiplasy versus convergence (Guerrero & Hahn, 2018), for identifying introgressed regions (Fang *et al.*, 2020), and testing hypotheses about adaptation (Martin *et al.*, 2019). Despite the development of advanced hierarchical models for inferring species trees, such methods have yet to be developed for spatially linked gene tree estimation.

Here we have demonstrated that the probability of incongruence described by the edge lengths of a species tree in coalescent units does not capture the expected spatial similarity of genealogies across chromosomes. Instead, in addition to the ratio of *t*_*g*_ to *N*_*e*_, which describes the probability of incongruence, it becomes necessary to consider the magnitudes of these parameters as well. This presents an interesting scenario: imagine a balanced tree with two clades where every edge has the same *t*_*c*_ edge lengths. In one clade these edges are composed of high *N*_*e*_ and *t*_*g*_ values, while in the other clade edges have low *N*_*e*_ and *t*_*g*_ values. Despite having the same probability of incongruence, the two clades would exhibit very different rates of change in their topology per unit length spatially across the genome. Unlike in our simulations, the rate of decay would likely not be uniform, and would covary more among some edges than others.

In theory, this expectation could be built into sliding window analyses based on a parameterized species tree inferred from unlinked loci. The simple approach that we implemented here, applying weights to alignment windows, is only a first step. A more appropriate direction to focus in the future would be to use species tree information to establish tree topology priors in a Bayesian context that could be used to improve local gene tree estimation by combining both the expected genome-wide distribution of genealogies as well as the expected similarity among neighboring genealogies. In contrast to treating recombination as a source of error for phylogenetic analyses, this direction of research aims to accommodate recombination as a source of historical information. There is no doubt that the MSC will continue to be extended to meet the needs introduced by new types of data, and the many questions that they inspire.

## Acknowledgments

We thank Lacey Knowles and Laura Kubatko for organizing the Estimating Species Trees book volumes and for inviting us to contribute. This work was supported by a NSF Graduate Research Fellowship to P. McKenzie and NSF grant DEB 1557059 to D. Eaton.

## References

Castillo-Ramirez, S., Liu, L., Pearl, D. & Edwards, S. (2010). Bayesian estimation of species trees: a practical guide to optimal sampling and analysis. In: Estimating species trees: practical and theoretical aspects. Wiley-Blackwell, pp. 15–33. 2

Degnan, J.H. & Rosenberg, N.A. (2009). Gene tree discordance, phylogenetic inference and the multi-species coalescent. Trends in Ecology & Evolution, 24, 332–340. 1

Eaton, D.A.R. (2020). Toytree: A minimalist tree visualization and manipulation library for Python. Methods in Ecology and Evolution, 11, 187–191. 2

Edwards, S. & Beerli, P. (2000). Perspective: Gene Divergence, Population Divergence, and the Variance in Coalescence Time in Phylogeographic Studies. Evolution, 54, 1839–1854. 1

Fang, B., Merilä, J., Matschiner, M. & Momigliano, P. (2020). Estimating uncertainty in divergence times among three-spined stickleback clades using the multispecies coalescent. Molecular Phylogenetics and Evolution, 142, 106646. 1, 7

Guerrero, R.F. & Hahn, M.W. (2018). Quantifying the risk of hemiplasy in phylogenetic inference. Proceedings of the National Academy of Sciences, 115, 12787–12792. 1, 7

Hudson, R.R. & Kaplan, N.L. (1988). The coalescent process in models with selection and recombination. Genetics, 120, 831–840. 1

Kelleher, J., Etheridge, A.M. & McVean, G. (2016). Efficient Coalescent Simulation and Genealogical Analysis for Large Sample Sizes. PLOS Computational Biology, 12, e1004842. 1, 2

Knowles, L.L. & Kubatko, L.S. (2011). Estimating Species Trees: Practical and Theoretical Aspects. John Wiley and Sons. 1

Maddison, W.P. (1997). Gene Trees in Species Trees. Systematic Biology, 46, 523–536. 1

Maddison, W.P. & Knowles, L.L. (2006). Inferring Phylogeny Despite Incomplete Lineage Sorting. Systematic Biology, 55, 21–30. 1

Martin, S.H. & Belleghem, S.M.V. (2017). Exploring Evolutionary Relationships Across the Genome Using Topology Weighting. Genetics, 206, 429–438. 2

Martin, S.H., Davey, J.W., Salazar, C. & Jiggins, C.D. (2019). Recombination rate variation shapes barriers to introgression across butterfly genomes. PLOS Biology, 17, e2006288. 7

McKenzie, P.F. & Eaton, D.A.R. (2020). ipcoal: An interactive Python package for simulating and analyzing genealogies and sequences on a species tree or network. bioRxiv, p. 2020.01.15.908236. 2

McVean, G.A. & Cardin, N.J. (2005). Approximating the coalescent with recombination. Philosophical Transactions of the Royal Society B: Biological Sciences, 360, 1387–1393. 1

Rasmussen, M.D. & Kellis, M. (2012). Unified modeling of gene duplication, loss, and coalescence using a locus tree. Genome Research, 22, 755–765. 1

Springer, M.S. & Gatesy, J. (2016). The gene tree delusion. Molecular Phylogenetics and Evolution, 94, 1–33. 3

Stamatakis, A. (2014). RAxML version 8: a tool for phylogenetic analysis and post-analysis of large phylogenies. Bioinformatics (Oxford, England), 30, 1312–1313. 6

Wang, J., Street, N.R., Park, E.J., Liu, J. & Ingvarsson, P.K. (2019). Evidence for widespread selection in shaping the genomic landscape during speciation of Populus. bioRxiv, p. 819219. 2

Yu, Y., Than, C., Degnan, J.H. & Nakhleh, L. (2011). Coalescent Histories on Phylogenetic Networks and Detection of Hybridization Despite Incomplete Lineage Sorting. Systematic Biology, 60, 138–149. 1

